# Oblique transmission, conformity, and preference in the evolution of altruism

**DOI:** 10.1101/2020.12.10.420513

**Authors:** Kaleda K. Denton, Yoav Ram, Marcus W. Feldman

## Abstract

The evolution of altruism is frequently studied using models of non-random assortment, including kin selection. In genetic kin selection models, under certain assumptions including additive costs and benefits, the criterion for altruism to invade a population is Hamilton’s rule. Deviations from Hamilton’s rule occur when vertical transmission has cultural and genetic components, or when costs and benefits are combined multiplicatively. Here, we include oblique and vertical cultural transmission and genetic transmission in four models—two forms of parent-to-offspring altruism, sibling-to-sibling altruism, and altruism between offspring that meet assortatively—under additive or multiplicative assumptions. Oblique transmission may be conformist (anti-conformist), where the probability that an individual acquires a more common cultural variant is greater (less) than its frequency. Inclusion of conformist or anti-conformist oblique transmission may reduce or increase the threshold for invasion by altruism relative to Hamilton’s rule. Thresholds for invasion by altruism are lower with anti-conformity than with conformity, and lower or the same with additive rather than multiplicative fitness components. Invasion by an allele that increases the preference for altruism does not depend on oblique phenotypic transmission, and with sibling-to-sibling altruism, this allele’s invasion threshold can be higher with additive rather than multiplicative fitnesses.

## 1 Introduction

Altruism occurs when an individual’s behaviour toward another causes the former to suffer a cost while the latter obtains a benefit from the former’s behaviour. Such behaviours are common in humans [1] and eusocial species of insects and mole-rats [2, 3, 4], and occur to varying degrees in many other taxa. How altruistic behaviours may evolve has intrigued biologists since Darwin [5], and has been the subject of a great deal of theoretical research [1, 6, 7, 8, 9, 10, 11, 12, 13].

Hamilton [7, 8] suggested a condition for the evolution of altruism that has since become known as Hamilton’s rule: genetically-determined altruism will invade a population of selfish individuals (non-altruists) if *γ < rβ*, where *γ* is the fitness cost to altruists, *β* is the fitness benefit to recipients of altruism, and *r* is the degree of relatedness between donors and recipients. This process of increasing the fitness of relatives was first called kin selection by Maynard Smith [14]. Hamilton’s rule can hold in population-genetic models that assume Hardy-Weinberg proportions among genotype frequencies after selection [15, 16, 17], known as inclusive fitness models, which are approximations of exact population-genetic models under weak selection [18, 19, 20].

In exact population-genetic models, where selection can be of any strength, Hamilton’s rule holds only under certain assumptions [9, 20, 21, 22, 23, 24]. Cavalli-Sforza and Feldman [9] modeled altruism as a genetic trait, controlled by a single locus with two alleles, and found that Hamilton’s rule could hold when costs and benefits combined additively but not multiplicatively (see also [21, 25]). Departures from Hamilton’s rule also occur if altruism is affected by more than one locus [24] or by multiple alleles at a single locus [22], or if genetic transmission is non-vertical (e.g., via associated microbes) [26, 27]. In Feldman et al. [28], altruism had a genetic component controlled by one allele at one locus, as well as a vertically-transmitted cultural component, and costs and benefits combined additively. They found that conditions for invasion by altruism differed from Hamilton’s rule, and conditions for invasion by an allele linked to the altruistic locus differed from Hamilton’s rule in the case of parent-to-offspring altruism.

A general form of Hamilton’s rule (known as HRG for Hamilton’s Rule General [29]) has been suggested, as well as different frameworks for understanding the genetic evolution of altruism, such as group selection [30, 31, 32, 33] and reciprocity [13, 34, 35]. HRG is derived from the Price [36] equation, and holds even if fitness combinations are non-additive and selection is strong [37, 38, 39]. However, HRG does not have predictive power (as *β* and *γ* incorporate the dependent variable, namely the change in average trait value over one generation) [29, 40, 41], and does not account for non-vertical transmission of altruism (but see [42]). Studies of altruism as a cultural trait under non-vertical transmission have commonly used the frameworks of group selection [43, 44, 45, 46, 47] or network reciprocity [48, 49, 50, 51], as opposed to kin selection (but see [28, 52]). These studies have reached opposing conclusions on whether conformity—a type of non-vertical transmission that has been defined in different ways [53] but generally refers to a tendency to adopt more common variants—facilitates [43, 44, 49, 50, 51] or hinders [45, 48] the evolution of altruism.

Here we build on the kin selection models of Feldman et al. [28], in which altruism had vertically-transmitted genetic and cultural components, and fitness components were additive. We incorporate oblique cultural transmission and explore cases of additive and multiplicative fitness components. Oblique transmission is assumed to be frequency-dependent in that the probability that offspring adopt a given variant is a function of its frequency in the parental generation. Two types of frequency-dependent transmission that we investigate are conformity and anti-conformity, using definitions from [44]: *under conformity (anti-conformity), the probability that an individual acquires a more common cultural variant is greater (less) than the variant’s frequency*.

Our analysis includes parent-to-offspring altruism (parental care) and sibling-to-sibling altruism, as in [28], and altruism between offspring that meet assortatively, as in [54], which approximates non-random interactions that may be due to kin selection, group selection, or other forms of population structure. Some definitions of altruism exclude parent-to-offspring altruism, but we use the definition from [10]: *altruists are individuals that sacrifice their fitness to benefit that of other individuals*. Altruists can be parents and recipients can be their offspring, and the reduction of parents’ fitness is in terms of future offspring not produced. Conditions are derived under which altruism, and an allele that increases the preference for altruism, can invade a population initially fixed on selfishness, or an allele corresponding to a lower preference for altruism, respectively.

## 2 Models

In a population of sexually reproducing haploids there are two possible phenotypes, altruism (phenotype 1) and selfishness (phenotype 2), and two genotypes, *A* and *a*, that affect the transmission of these phenotypes. The four phenogenotypes, *A*_1_, *A*_2_, *a*_1_, *a*_2_, have frequencies *u*_1_, *u*_2_, *u*_3_, *u*_4_, respectively, in the parental generation, *ũ*_1_, *ũ*_2_, *ũ*_3_, *ũ*_4_, in the offspring generation after transmission, and 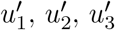, and 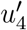 after selection, which follows transmission.

### 2.1 Transmission

Transmission is vertical with probability *ρ* and oblique with probability 1 *− ρ*. If vertical transmission occurs, as in [28], one of the two parents is randomly chosen to be the “transmitting parent.” An offspring initially acquires its transmitting parent’s phenotype and then adjusts its phenotype based on its preference, which is affected by genotypes *A* and *a*; thus, 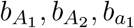, and 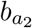 represent phenogenotypes’ preferences for altruism. An offspring with initial phenogenotype *i* becomes altruistic with probability *b*_*i*_ and selfish with probability 1 *− b*_*i*_ (Figure 1). Purely genetic transmission entails that 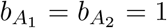 and 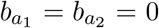, in which case individuals with allele *A* are altruists and those with *a* are selfish.

**Figure 1:**
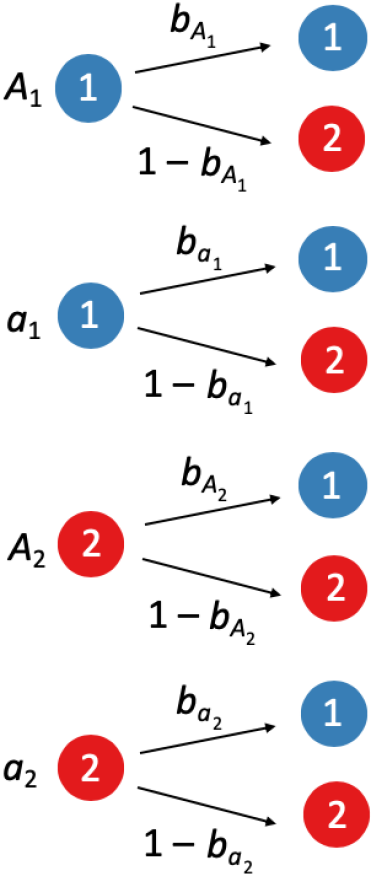
Preference parameters, *b*_*i*_. In the left column, four offspring have initially inherited either phenotype 1, altruism (in blue) or phenotype 2, selfishness (in red) due to vertical or oblique transmission. They have also inherited either allele *A* or *a*, and their phenogenotypes are shown to their left. The right column depicts these offspring either retaining their original phenotype or switching to the other phenotype, with probabilities given beside the respective arrows. If the initial phenogenotype was *i*, the probability that the final phenotype is 1 is *b*_*i*_ and the probability the final phenotype is 2 is 1 *− b*_*i*_.

If transmission is oblique, the probability that an offspring initially acquires altruism (before exercising its preference, *b*_*i*_) is *f* (*u*_1_ + *u*_3_), and the probability that it initially acquires selfishness is *f* (*u*_2_ + *u*_4_) = 1 *− f* (*u*_1_ + *u*_3_). *f* (*x*) may take any form of frequency-dependent transmission with *f* (0) = 0, *f* (1) = 1, and for a small frequency, *ε* (because we focus on initial increase), *f* (*ε*) = *cε*+*O*(*ε*^2^), where *c* is a positive constant. In the conformity model of Boyd and Richerson [44], if *n* is the number of cultural role models that an offspring randomly samples from the parental generation, then 0 *< c <* 1 with conformity and 1 *< c < n* with anti-conformity (see electronic supplementary material [hereafter, ESM] 1 and Eq. (29) of [55]). Ultimately, under oblique transmission, the probability that an offspring of genotype *A* (*a*) becomes altruistic is *S* (*U*) and the probability that it becomes selfish is *T* (*V*), where

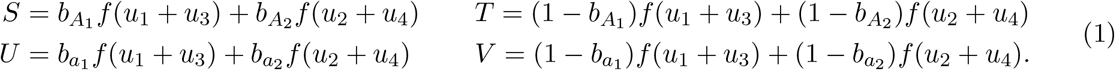

Let *W* (*Y*) be the probability that the transmitting parent is altruistic and the offspring has allele *A* (*a*), and *X* (*Z*) be the probability that the transmitting parent is selfish and the offspring has allele *A* (*a*), as in ESM 2.

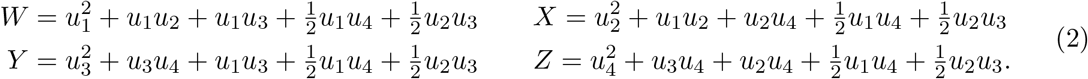

After vertical and oblique transmission, the frequencies of *A*_1_, *A*_2_, *a*_1_, *a*_2_ are, respectively,

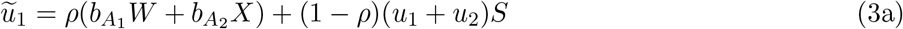

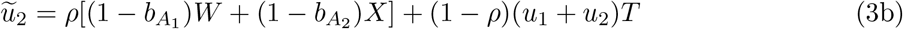

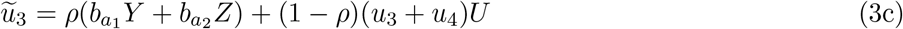

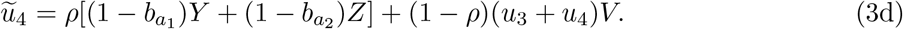

### 2.2 Selection

Selection follows transmission. Altruistic offspring suffer a fitness cost *γ* (0 *< γ <* 1) whereas selfish offspring do not. Offspring may also receive a fitness benefit *β* (0 *< β*) from either a parent (Models I and II), sibling (Model III), or member of the offspring generation (Model IV) that is altruistic.

#### 2.2.1 Additive Models

Let *y* refer to the potential donor (either a parent, a sibling, or another member of the offspring generation), which, if it is an altruist, donates a fitness benefit to the recipient. Then

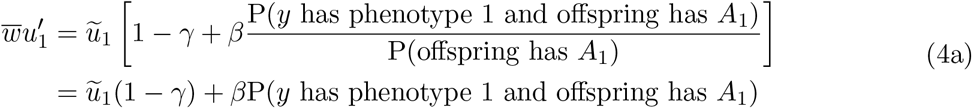

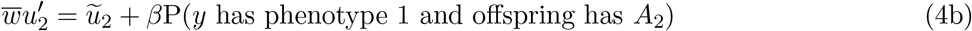

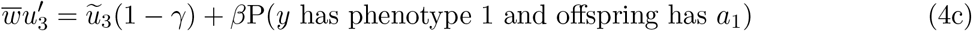

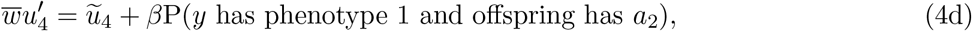

where the normalizer, 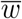, is the sum of the right-hand sides, which we refer to as mean fitness, and ũ_1_, *ũ*_2_, *ũ*_3_, *ũ*_4_ are given by (3).

In Model I, the potential donor, *y*, is a randomly-selected parent. The calculation of P(*y* has phenotype 1 and offspring has *A*_1_) is in ESM 2, and the corresponding parts of equations (4b-d) are calculated similarly. Table S2.1 shows the possible mating pairs and corresponding offspring phenogenotypes and Table S2.2 includes the probability that the parent that is selected to benefit the offspring is altruistic (ESM 2). The recursions after transmission and selection in Model I are

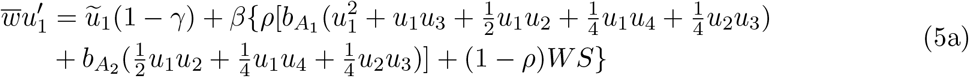

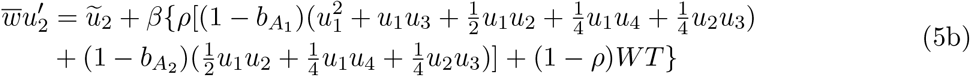

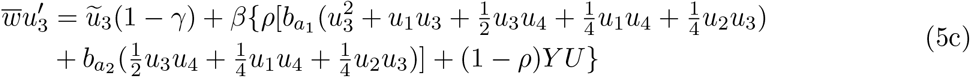

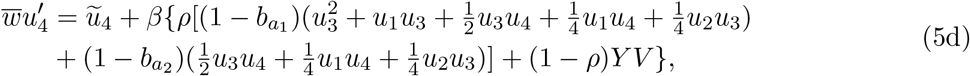

where 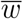 is the mean fitness, *S, T, U, V* are given in (1) and *W, X, Y, Z* are given in (2).

In Model II, the potential donor, *y*, is the transmitting parent. The difference between Models I and II is illustrated in Figure S3.1 (ESM 3); the oblique terms are the same as in Model I and the vertical terms change in the same way as in Model II of [28]:

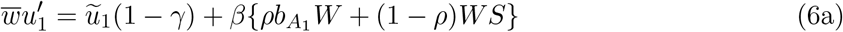

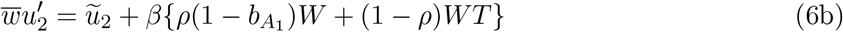

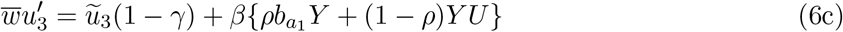

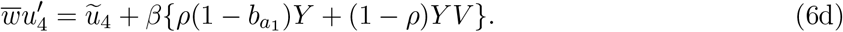

In Model III, the potential donor, *y*, is a sibling, and the transmitting parent is the same for the two siblings (i.e., uniparental transmission). ESM 2 shows the calculation of P(*y* has phenotype 1 and recipient has *A*_1_), and the corresponding parts of (4b-d) are calculated similarly. Note that because both siblings’ phenotypes depend on preferences *b*_*i*_, there are terms with 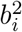, whereas *b*_*i*_ only appears to the first power in the parent-to-offspring altruism models. The recursion for *A*_1_ in Model III is below and the other recursions are in ESM 4, equations (S4.1).

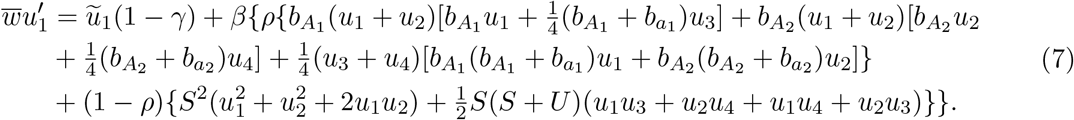

In Model IV, the potential donor, *y*, is another offspring. Following the assortative meeting model of Eshel and Cavalli-Sforza [54], let 0 *< m <* 1 be the probability that an individual non-randomly encounters another of the same phenotype, and 1 *− m* be the probability of random encounters. The probability that the potential donor, *y*, is an altruist (after transmission) is ũ _1_ +ũ_3_, and for each altruist, the probability of encountering another altruist is *m* + (1 *− m*)(ũ_1_ + ũ_3_), as in [54]. For a selfish individual, the probability of encountering an altruist is (1 *− m*)(ũ _1_ + ũ _3_). The recursions are

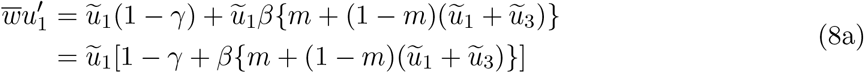

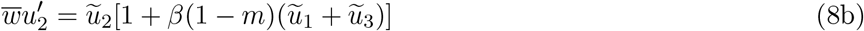

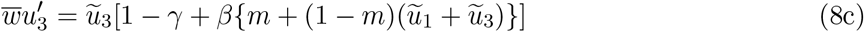

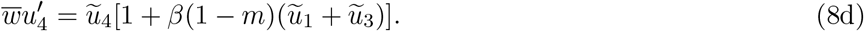

#### 2.2.2 Multiplicative Models

If costs and benefits combine multiplicatively, then

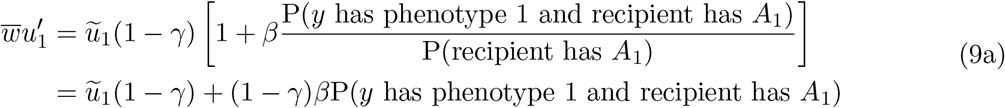

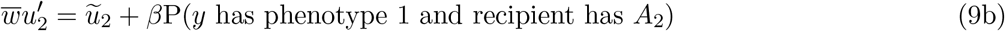

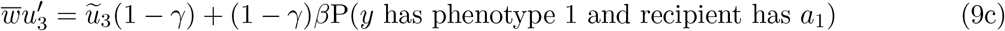

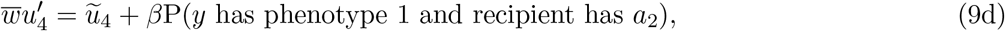

where 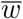 is the mean fitness. Recursions (9b) and (9d) are the same as additive recursions (4b) and (4d), respectively, whereas the recursions for altruistic phenogenotypes differ; in (9a) and (9c) (1 *− γ*)*β* appears where *β* appeared in (4a) and (4c).

## 3 Conditions for Invasion by Altruism

### 3.1 Invasion by Altruism in the Additive Case

Suppose that the population is initially fixed on selfishness, in which case selfish individuals cannot become altruistic (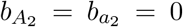; see ESM 5). There is no selection on *A* versus *a*, and the initial equilibrium is a point on the neutral curve (*u*_1_, *u*_2_, *u*_3_, *u*_4_) = (0, *û*_2_, 0, *û*_4_). Near one of these equilibria, altruistic phenogenotypes *A*_1_ and *a*_1_ are introduced at small frequencies, denoted by *u*_1_ = *ε*_1_ and *u*_3_ = *ε*_3_. We assume that the preference for altruism (Figure 1) is genotype-independent with 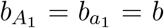 as in [28]; otherwise, invasion conditions become very difficult to calculate and interpret (e.g., see Eq. (S5.1) in ESM 5). For all models, the sum of the linearized recursions for frequencies of *A*_1_ and *a*_1_, respectively, is (ESM 5)

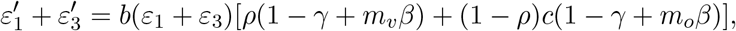

where *m*_*o*_ and *m*_*v*_ vary depending on the model and are given in Result 1 below. Altruism increases if 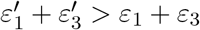, expressed as Result 1.

**Result 1. In additive models, the condition for invasion by altruism of a population that is initially all selfish is**

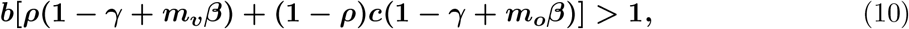

**where *m***_***v***_ **and *m***_***o***_ **are assortment parameters for vertical and oblique transmission, respectively**.

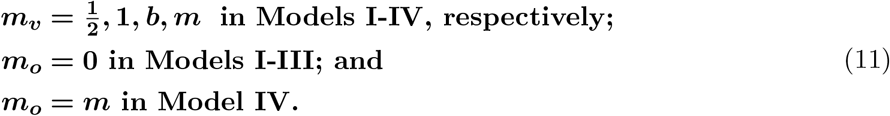

Inequality (10) can be rearranged to 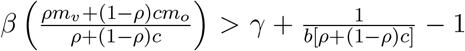, where 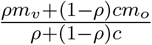 takes the place of relatedness (*r*) in Hamilton’s rule and assortment (*m*) in *mβ > γ*. The term 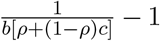 is the adjustment due to transmission bias, which raises the threshold for invasion if it is positive and lowers the threshold if it is negative. This term is negative if there is sufficiently strong anti-conformist oblique transmission (i.e., 0 *≤ ρ <* 1 and 1 *< c < n*).

By (10), increasing *c* and increasing *b* (the preference for altruism) facilitates invasion by altruism. Examples for Model I are in Figure 2, and examples for all models are in ESM 6 (code is available at https://github.com/kaleda/altruism-conformity). The relationship between inequality (10) and *ρ* is more complicated. If *ρ* = 1 inequality (10) becomes

**Figure 2:**
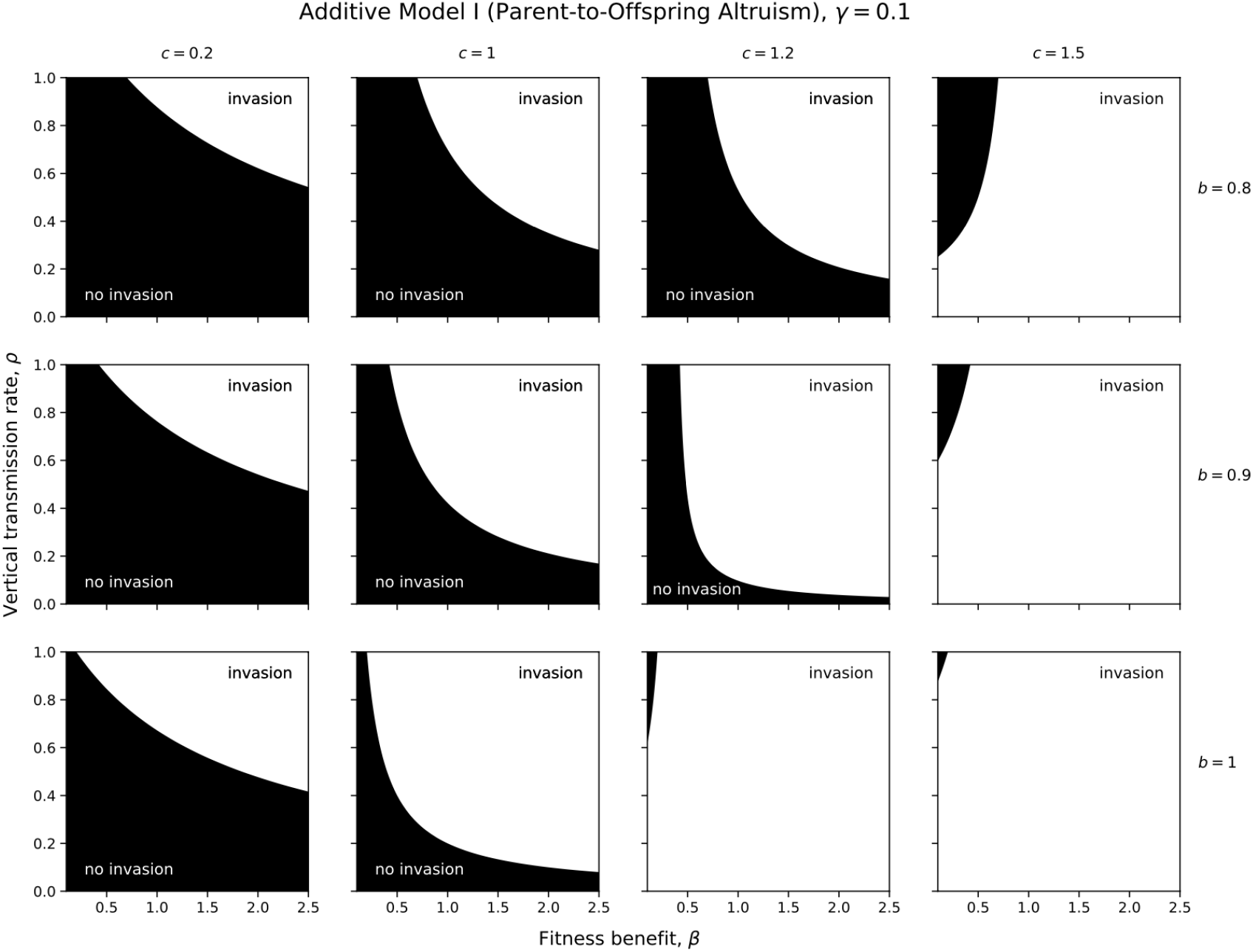
Conditions under which altruism invades in additive Model I (inequality 10), where the cost of altruism (*γ*) is fixed at 0.1. In light areas, (10) holds and altruism invades; in dark areas, it does not. In the far left column there is conformity, in the second column from the left there is random copying, and the two right-most columns there is anti-conformity. The top row includes a weaker preference for altruism than the middle row, and the bottom row includes the strongest possible preference for altruism. Recall that 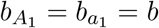 in deriving inequality (10).

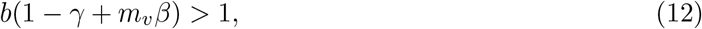

which gives the same results as in [28] for Models I-III. If *ρ* = 0, inequality (10) becomes

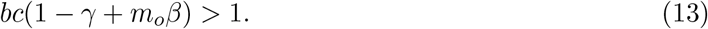

If the left-hand side of (12) is greater than the left-hand side of (13), increasing *ρ* (vertical transmission) facilitates invasion (by (10)). If the opposite is true, decreasing *ρ* facilitates invasion.

#### Remark 1.

*(i) In all models, m*_*v*_ *≥ m*_*o*_; *therefore, with conformity* (0 *< c <* 1), *increasing ρ facilitates invasion by altruism. (ii) In Model IV, m*_*v*_ = *m*_*o*_, *and with anti-conformity* (1 *< c < n*) *decreasing ρ facilitates invasion by altruism*.

Inequality (10) can be compared to the condition for invasion by altruism under purely genetic transmission, namely Hamilton’s rule, 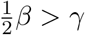, for Models I-III [28] and *mβ > γ* for Model IV [54]. We rearrange (10) to isolate 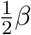 (in Models I-III) or *mβ* (in Model IV) on the left-hand side, and then determine when the right-hand side is less than *γ*, producing a lower threshold for invasion than in the case of purely genetic transmission. We find

**Result 2. In additive models, the threshold for invasion by altruism in (10) is lower than the threshold for invasion with purely genetic transmission if**

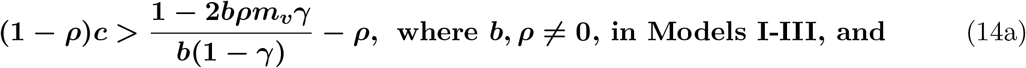

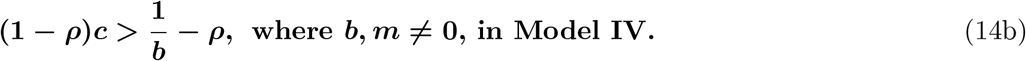

Recall that 0 *< γ <* 1, 0 *< c <* 1 with conformity, and 1 *< c < n* with anti-conformity. In Models I and IV, inequalities (14a,b), respectively, cannot hold if *ρ* = 1 or if *ρ <* 1 and oblique transmission is conformist. However, with anti-conformity, if *c* is large enough (due to a large enough number of role models, *n*), inequalities (14a,b) may hold. In Models II and III, inequality (14a) can hold with either *ρ* = 1 or 0 *< ρ <* 1, and in the latter case, with either conformity or anti-conformity.

We also compare invasion condition (10) under a mixture of vertical and oblique transmission (0 *< ρ <* 1; hereafter, “mixed” transmission) to the invasion condition with completely vertical transmission (*ρ* = 1) given in (12). We isolate *β* on the left-hand side of inequality (10) in the cases of mixed transmission and completely vertical transmission, and determine when the right-hand side for the former can become smaller than that for the latter, meaning that the threshold for invasion is lower with mixed than with completely vertical transmission.

**Result 3. In additive models, the threshold for invasion by altruism is lower with mixed transmission (0 *< ρ <* 1) than completely vertical transmission (*ρ* = 1) if**

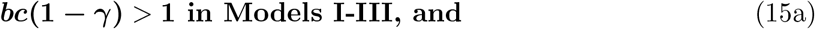

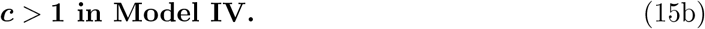

Inequalities (15a,b) do not hold with conformity, in accordance with Remark 1 (i). Inequality (15a) may hold with anti-conformity (1 *< c < n*) if *c* is sufficiently large, and (15b) always holds with anti-conformity, in accordance with Remark 1 (ii).

#### Remark 2.

*In Models I-III with completely oblique transmission* (*ρ* = 0), *from inequality (13), invasion by altruism occurs if and only if (15a) holds*. Therefore, in Models I-III, if the threshold for invasion is higher with mixed transmission than with completely vertical transmission, invasion is impossible with completely oblique transmission.

Finally, the invasion threshold with mixed transmission is compared to that with completely oblique transmission (*ρ* = 0) given in (13), except that rather than isolating *β* on one side of the two inequalities, *γ* is isolated (because *β* does not appear in inequality (13) for Models I-III).

**Result 4. In additive models, the threshold for invasion by altruism is lower with mixed transmission (0 *< ρ <* 1) than completely oblique transmission (*ρ* = 0) if**

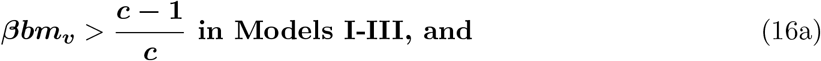

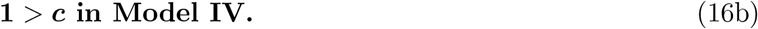

Inequalities (16a,b) hold with conformity, in accordance with Remark 1 (i). With anti-conformity, (16a) may hold, but (16b) never holds, in accordance with Remark 1 (ii).

### 3.2 Invasion by Altruism in the Multiplicative Case

Conditions for invasion by altruism are derived from recursions in *u*_1_ and *u*_3_, both of which have *β* replaced with (1 *− γ*)*β* in the multiplicative model. Therefore, Result 5 differs by Result 1 in that *β* is replaced by (1 *− γ*)*β* wherever it appears.

**Result 5. In multiplicative models, the condition for invasion by altruism of a population that is initially all selfish is**

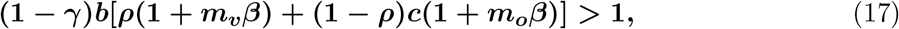

**where *m***_***v***_ **and *m***_***o***_ **for each model are given in (11)**. Due to the reduction in the term multiplying *β* in the multiplicative case, the threshold for invasion by altruism is higher in the multiplicative than additive case in all models except when *ρ* = 0 in Models I-III, in which cases these thresholds are the same. It can be shown that Remarks 1 and 2 and Result 3 apply in multiplicative as well as additive cases. Result 2 applies in the multiplicative case if the genetic model being compared is also multiplicative.

## 4 Conditions for Invasion by Allele *A*

### 4.1 Invasion by Allele *A* in the Additive Case

Here, we find conditions for invasion by allele *A* of a population initially fixed on *a*. To simplify this analysis, as in [28], we assume that 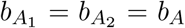 and 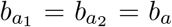, where *b*_*A*_ (*b*_*a*_) is the preference of carriers of allele *A* (*a*) for altruism. Because the probability that an individual is an altruist does not depend on which phenotype it acquired initially, only its allele, the extent (1 *− ρ*) and form (*c*) of oblique transmission does not appear in recursions. Therefore, invasion conditions for additive Models I-III are the same as those in [28], where *ρ* = 1 was assumed (proof in ESM 7).

In additive Models I and II, there is no straightforward relationship between Hamilton’s rule, the preference difference *b*_*A*_ *− b*_*a*_, and the condition for invasion [28]. In additive Model III, allele *A* invades if 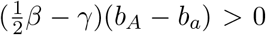; therefore, an allele that increases the preference for altruism (*b*_*A*_ *> b*_*a*_) invades if and only if Hamilton’s rule holds [28]. Similarly, for Model IV (ESM 7),

**Result 6. In additive Model IV (assortative meeting), the condition for invasion by allele *A* of a population that is initially fixed on allele *a* is**

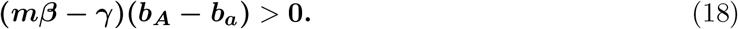

If *b*_*A*_ *> b*_*a*_, (18) reduces to the condition for invasion by altruism with genetic transmission, *mβ > γ*.

### 4.2 Invasion by Allele *A* in the Multiplicative Case

The allele invasion analyses for Models I-IV (ESM 7) are repeated, but with *β*(1 *− γ*) replacing *β* in the recursions for *u*_1_ and *u*_3_. These analyses are in ESM 8, and the results are below. Recall 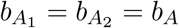 and 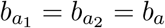.

**Result 7. In multiplicative Models I and II (parent-to-offspring altruism), the condition for invasion by allele *A* of a population that is initially fixed on allele *a* is**

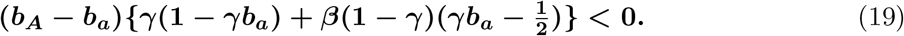

If *b*_*A*_ *> b*_*a*_, invasion by allele *A* occurs if 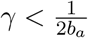 and 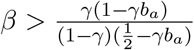. Comparing invasion condition (19) in the multiplicative model to the corresponding invasion condition in the additive model is not straightforward because the latter involves the root of a quadratic (ESM 7), so we conducted a numerical analysis (code is available at https://github.com/kaleda/altruism-conformity). For values of *β, γ, b*_*a*_, and *b*_*A*_ separated by 0.01 (0.01 *−* 10^*−*5^ at bounds) and with *β ≤* 10 and *b*_*A*_ *> b*_*a*_, we found no instances of invasion in the multiplicative, but not additive, model. Approximately 31.4% of cases showed invasion in the additive, but not multiplicative, model.

**Result 8. In multiplicative Model III (sibling-to-sibling altruism), the condition for invasion by allele *A* of a population that is initially fixed on allele *a* is**

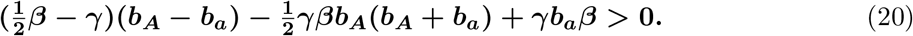

It can be shown (ESM 8) that the threshold for invasion by allele *A* in Model III is lower in the multiplicative than additive case if 2*b*_*a*_ *> b*_*A*_(*b*_*A*_ + *b*_*a*_), which always holds if *b*_*A*_ *< b*_*a*_.

**Result 9. In multiplicative Model IV (assortative meeting), the condition for invasion by allele *A* of a population that is initially fixed on allele *a* is**

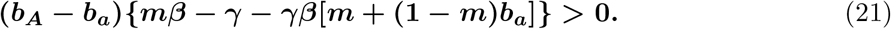

If *b*_*A*_ *> b*_*a*_, a necessary condition for invasion by *A* is 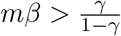. Compared to (18) for the additive case, the invasion threshold in the multiplicative case (21) is lower if and only if *b*_*A*_ *< b*_*a*_ (ESM 8).

## 5 Discussion and Conclusions

In our models, individuals are haploid and sexually reproducing, transmission is vertical with probability *ρ* or oblique with probability 1 *−ρ*, and alleles *A* and *a* influence the tendency to become altruistic through the preference parameters 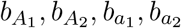 (Figure 1). Oblique transmission may be conformist or anti-conformist depending on the bias parameter *c*. Following Boyd and Richerson [44], 0 *< c <* 1 with conformity and 1 *< c < n* with anti-conformity, where *n* is the number of cultural role models. We explore two kinds of parent-to-offspring altruism (Models I and II), sibling-to-sibling altruism (Model III), and altruism between offspring that meet assortatively (Model IV).

To study invasion by altruism, we assume that the population is initially comprised of selfish individuals (*A*_2_ and *a*_2_) that cannot become altruistic 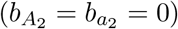, and that the preference for altruism is genotype-independent 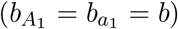. The threshold for invasion by altruism is lower in the additive case than the multiplicative case for all models except when *ρ* = 0 in Models I-III, where these thresholds are the same. In all models, the threshold for invasion by altruism decreases with increasing preference for altruism, *b*, and increasing tendency to anti-conform, *c* (Figure 2 and ESM 6). When a new phenotype appears, conformity acts against its adoption because it is rare, whereas anti-conformity facilitates its spread.

It is important to note that the roles of conformity and anti-conformity change as the frequency of altruism changes, and conformity can facilitate the increase of altruism if it reaches a sufficiently high frequency. Moreover, we assumed a random choice of role models, whereas if the population were subdivided into groups [43, 44, 45, 47] or connected in networks [48, 49, 50, 51], the roles of conformity and anti-conformity in the evolution of altruism may differ. In a group selection model for the evolution of altruism as a cultural trait, Molleman et al. [47] showed that if altruism had initially reached fixation in a single sub-group (e.g., due to stochastic effects), conformity may favor its increase in the meta-population, but if altruism arose due to rare mutations in the initially selfish population, conformity prevented its invasion.

We compared the threshold for invasion by altruism with cultural transmission to that with purely genetic transmission. Although we performed this analysis in the additive case, our results also hold when comparing the multiplicative versions of the two invasion conditions. In all models, the threshold for invasion can be lower with cultural transmission if there is mixed transmission (0 *< ρ <* 1) and oblique transmission is anti-conformist. In Models II and III, the threshold can also be lower with cultural transmission if cultural transmission is completely vertical (*ρ* = 1) or mixed (0 *< ρ <* 1) with conformist oblique transmission. Thus, although the threshold for invasion by altruism is lower with anti-conformity than with conformity, in Models II and III the invasion threshold can be lower with conformity than with purely genetic transmission.

We also compared the threshold for invasion by altruism with mixed transmission to the threshold for invasion with completely vertical or completely oblique transmission. These analyses were performed in the additive case, but the following findings also hold in the multiplicative case. With conformity, increasing the extent of vertical transmission, *ρ*, lowers the threshold for invasion in all models. In Model IV with anti-conformity, decreasing *ρ* lowers the threshold for invasion. The relationship between *ρ* and invasion thresholds in Models I-III with anti-conformity is more complicated. Interestingly, in Models I-III, the threshold for invasion is lower with mixed transmission than with completely vertical transmission if *bc*(1 *− γ*) *>* 1, which is also the condition for invasion with completely oblique transmission (*ρ* = 0). This condition does not involve the fitness benefit, *β*; if altruism invades under completely oblique transmission, it does so not because of the fitness benefits it provides, but because of transmission biases. Similarly, Ram et al. [56] show that oblique transmission can prevent disfavored phenotypes from extinction by enabling the transmission of these phenotypes independently of reproduction.

To find conditions under which allele *A* invades a population fixed on *a*, we set 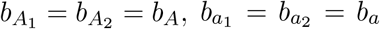, where *b*_*A*_ (*b*_*a*_) is the preference for altruism of carriers of allele *A* (*a*). Invasion conditions for additive Models I-III are identical to those in [28], where *ρ* = 1 was assumed.

In additive Models I and II, there is no straightforward relationship between the invasion condition, Hamilton’s rule, and the preference difference (*b*_*A*_ *− b*_*a*_). In additive Models III and IV, if *b*_*A*_ *> b*_*a*_ (*A* produces a greater preference for altruism than *a*), invasion occurs if 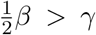 or *mβ > γ*, respectively, which are the invasion conditions under purely genetic transmission [28, 54].

The threshold for invasion by an allele that increases the preference for altruism (*b*_*A*_ *> b*_*a*_) is higher in the multiplicative case than the additive case for Model IV. Similarly, in Models I and II, we found many examples of invasion in the additive, but not multiplicative, case, but not vice versa. However, in Model III, the threshold for invasion by *A* with *b*_*A*_ *> b*_*a*_ may be lower in the multiplicative case than the additive case.

In these models, the genotype that determined the preference for altruism included one allele at one locus. However, an individual’s preference for altruism may be affected by more than one locus (as in [24]) or more than one allele at a given locus (as in [22]). It would be interesting to determine how a more complex relationship between genotype and preference for altruism affects the results of these models. In addition, in these models, encounters occurred between parents and offspring, between siblings, or between individuals that assorted by phenotype. Results would likely differ if encounters occurred between other kinds of relatives or if assortment were incorporated differently (see [57, 58] for a general treatment of non-random encounters).

In conclusion, when altruism is culturally transmitted, conditions for its invasion differ from those under purely genetic transmission. The threshold for invasion by altruism is lower with anti-conformity than with conformity, and in some models, can be lower with anti-conformity or conformity compared to purely genetic transmission. Incorporating additive rather than multiplicative fitness components produces a lower invasion threshold in all models except Models I-III with entirely oblique transmission, where these thresholds are identical. For an allele, *A*, that produces a greater preference for altruism than the resident allele, *a*, invasion conditions do not depend on the extent of oblique transmission or conformity. Invasion conditions for additive Models III and IV are the same as those under purely genetic transmission, whereas invasion conditions for additive Models I and II differ from Hamilton’s rule. In all models, the threshold for invasion by *A* may be lower under additive than multiplicative fitness combinations, and in Model III it can also be lower under multiplicative than additive combinations.

## Acknowledgements

This research was supported by the Stanford Center for Computational, Evolutionary and Human Genomics (KKD, MWF), Morrison Institute for Population and Research Studies at Stanford University (KKD, MWF), Israel Science Foundation 552/19 (YR), and Minerva Stiftung Center for Lab Evolution (YR).

## Author Contributions

KKD and MWF designed the research; KKD, YR, and MWF performed the research and wrote the paper.

